# standR: a Bioconductor package for analysing transcriptomic Nanostring GeoMx DSP data

**DOI:** 10.1101/2023.04.23.538017

**Authors:** Ning Liu, Dharmesh D. Bhuva, Ahmed Mohamed, Micah Bokelund, Arutha Kulasinghe, Chin Wee Tan, Melissa J Davis

## Abstract

To gain a better understanding of the complexity of gene expression in normal and diseased tissues it is important to account for the spatial context and identity of cell *in situ*. State-of-the-art spatial profiling technologies, such as the Nanostring GeoMx Digital Spatial Profiler (DSP), now allow quantitative spatially resolved measurement of the transcriptome in tissues. However, the bioinformatics pipelines currently used to analyse GeoMx data often fail to successfully account for the technical variability within the data and the complexity of experimental designs, thus limiting the accuracy and reliability of subsequent analysis. Carefully designed quality control workflows, that include in-depth experiment-specific investigations into technical variation and appropriate adjustment for such variation can address this issue. Here we present *standR*, a R/Bioconductor package that enables an end-to-end analysis of GeoMx DSP data. With four case studies from previously published experiments, we demonstrate how the standR workflow can enhance the statistical power of GeoMx DSP data analysis and how application of standR enables scientists to develop in-depth insights into the biology of interest.

## INTRODUCTION

Quantitative gene expression analysis of disease systems using technologies such as bulk RNA-seq have led to many biomarker discoveries and mechanistic insights through application of differential expression, transcriptional network and pathway analysis methods (1,2). Single-cell RNA sequencing added further resolution to transcriptomics studies by enabling the investigation of the whole transcriptome at a single cell level, fueling the identification of many novel cell states (3,4). New generation spatial molecular measurement platforms that incorporate spatial information with existing imaging and sequencing technologies allows in-depth and fine-grained analyses, such as cell-cell interactions, cellular neighbourhood analysis and cell type deconvolution (5,6). Further, these technologies enable spatially resolved questions, such as the identification of differential expression between different parts of a tumour, between tissues with and without a particular cellular infiltrate, or of tissues adjacent to and distant from certain anatomical features.

Amongst the spatial platforms, Nanostring’s GeoMx Digital Spatial Profiler (DSP) (7) is one of the more robust platforms for Formal-Fixed Paraffin-Embedded (FFPE) tissues (8), providing regions of interest (ROIs) level-selection methods, with ROIs ranging from tens to hundreds of cells. The FFPE compatibility allows the GeoMx DSP to be applicable to clinical and pathological investigations using banked FFPE archival tissues, thus enabling retrospective clinical cohort studies. However, the generation of DSP data includes placing tissue samples on glass slides, where different slides may introduce technical variations to the data, becoming a source of batch effects, which can lead to false discovery (9). Besides, sampling biases, such as unbalanced cell count and segment size, could be present due to the nature of randomness of FFPE materials or tissue segments from patients. Taking these factors together with other technical variations that are commonly seen in bulk or mini-bulk RNA-seq experiments (*e.g.* sequencing errors and sequencing depth or library size (10), it is necessary to perform data quality control (QC) and filtering and appropriate normalization or batch correction when analysing GeoMx DSP data. Moreover, it has been long-established that linear-based method such as Limma (11,12) and edgeR (13) are more appropriate for carrying out differential expression (DE) analysis compared to traditional T-test (14), especially for datasets with limited sample size. Methods must also correctly account for the complexity of experimental designs in spatial data, where multiple samples may be taken from one patient, or from adjacent regions in a single tissue. Taken together, it is essential to construct a computational workflow that can carry out comprehensive QC, data normalization and can be compatible with complex experimental designs and sophisticated DE methods.

Based on our literature review of publications with GeoMx DSP transcriptome datasets from 2020 to July 2022 (Figure 1A), the current most generic approaches rely heavily on the default quality control of the platform as well as standard paired T-tests (Figure 1D), which may be inadequate to handle the complexity of the experiment. To this end, we have developed a Bioconductor R package *standR* (**S**patial **t**ranscriptomics **a**nalyses a**n**d **d**ecoding in **R**) to assist the QC, normalization and batch correction, differential expression analysis, and downstream analysis of Nanostring GeoMx transcriptomics data. Here we introduce *standR* and describe the package’s workflow and utility in analysing GeoMx DSP datasets. We have also performed a comprehensive comparison of *standR* with the current generic GeoMx DSP workflow on four publicly available GeoMx datasets.

**Figure 1.**
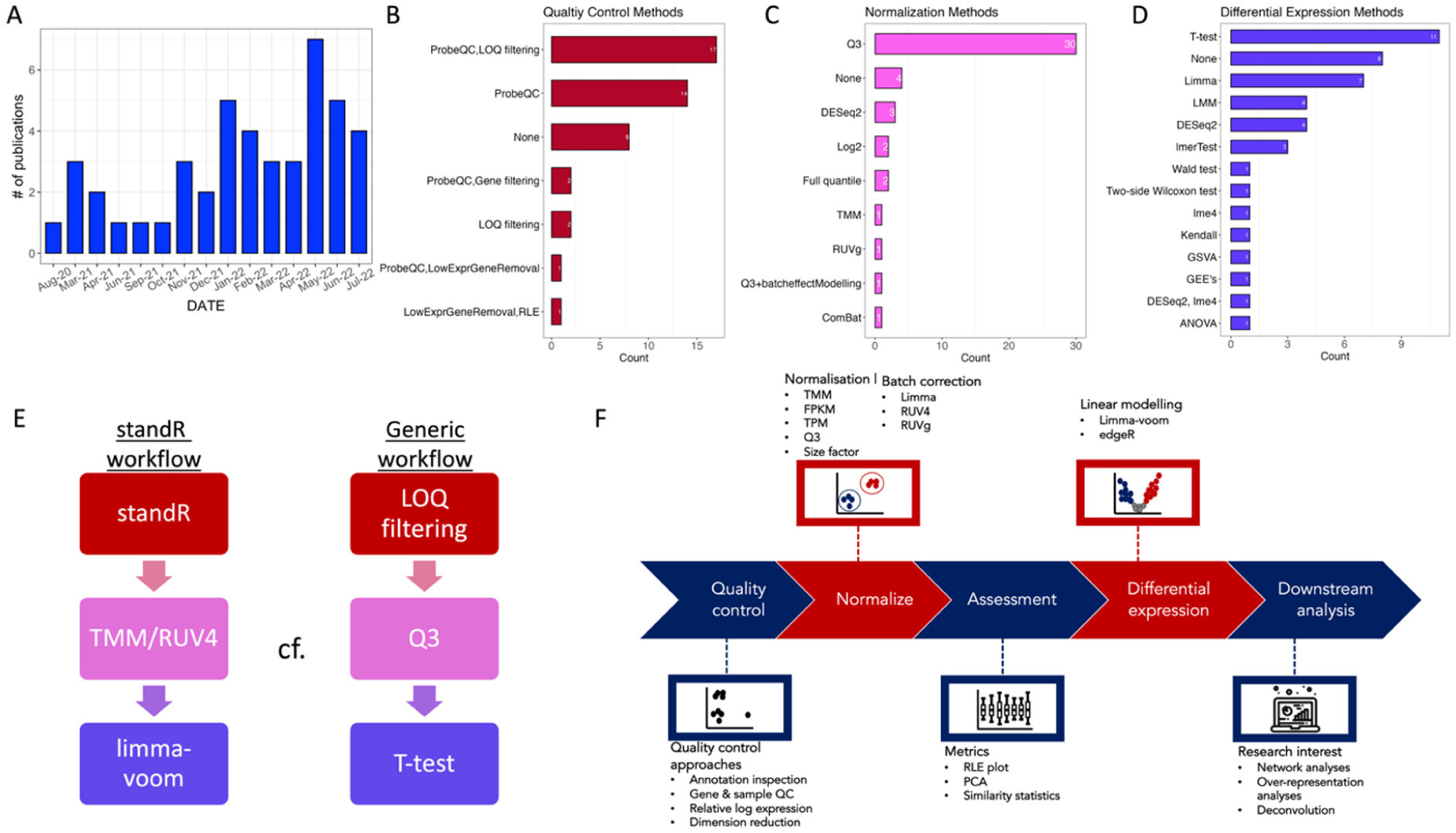
Literature Review and the standR workflow. A: Bar plot shows the increasing trend of publications with Nanostring GeoMx DSP datasets. B-D: Bar plots show the preferential choices of QC, normalization and differential expression methods in publications. E: Diagram demonstrates the comparison between standR and generic workflows. F: Flow diagram shows the standR workflow.

## MATERIAL AND METHODS

### Nanostring GeoMx DSP data pre-processing

The Nanostring GeoMx datasets used in this study are publicly available and were downloaded from Nanostring’s Spatial Organ Atlas (https://nanostring.com/products/geomx-digital-spatial-profiler/spatial-organ-atlas/). Initial data processing and sample-based QC were conducted using *standR*, any ROIs that were assigned with QC flags indicating low qualities (including “Low Percent Aligned Reads”, “Low Percent Stitched Reads”, “Low Surface Area”, “Low Nuclei Count”) were excluded from further analysis.

### Quality control

For the gene filtering, the generic workflow removes genes with raw count smaller than the ROI-specific limitation of quantification (LOQ) in all ROIs, while the standR workflow removes genes with logCPM (log counts per million) count smaller than a calculated threshold in 90% of the ROIs. The threshold is calculated based on the median and mean of library size and a minimal count (default is 5) within the standR packages’s *addPerROIQC* function. QC was assessed by visualizing the mean-variance distribution of genes (Figure 2B), which was generated using *voom* from the *limma* R package with the linear modelling equation: *model.matrix(∼0 + TissueGroups)*.

**Figure 2.**
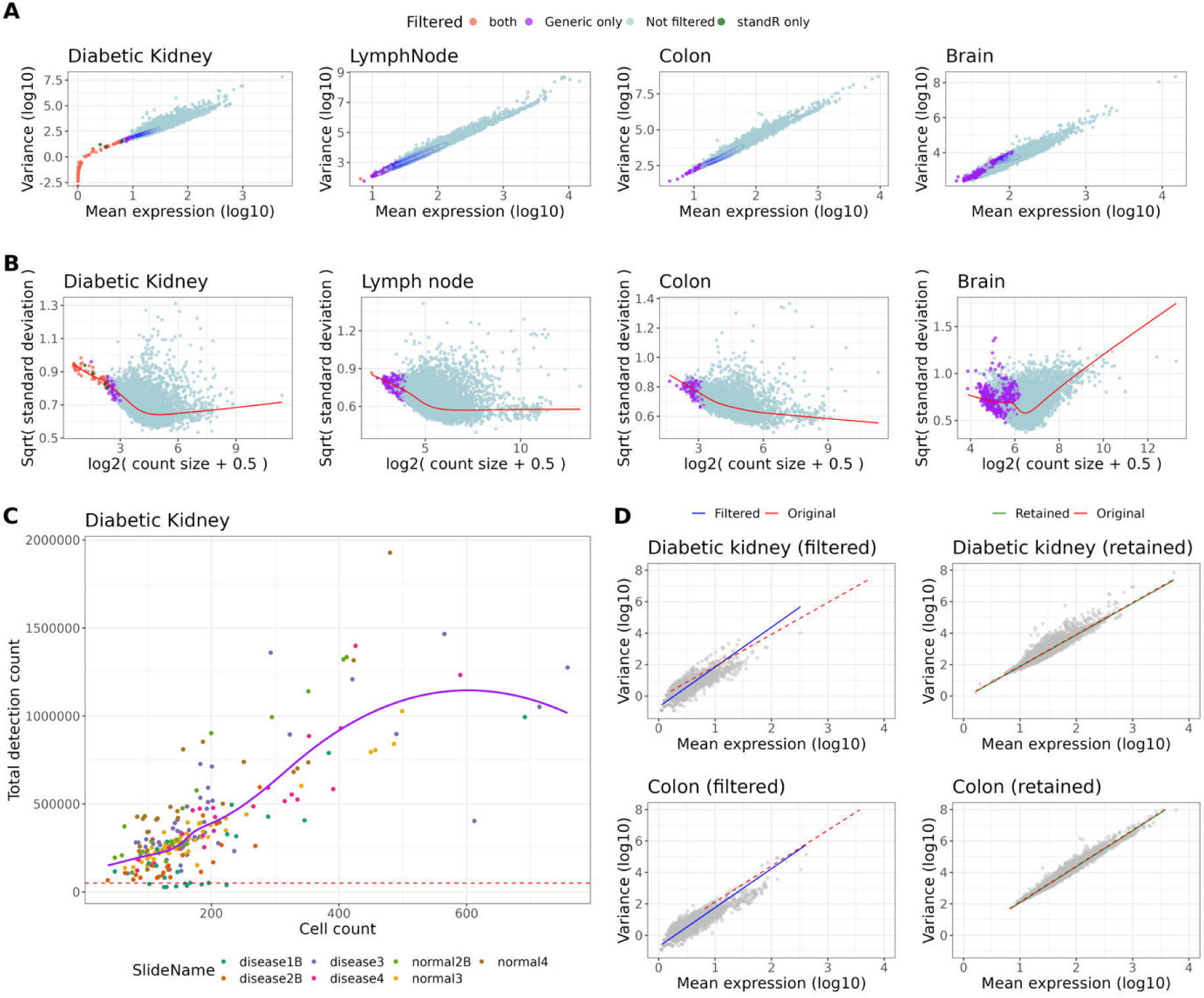
Gene and sample filtering by standR retains tissue relevant genes while removing low quality samples. Gene filtering (A&B), sample filtering (C&D). (A) Mean expression-variance of the genes across all the samples in each GeoMx dataset, colors denote if genes are removed by the gene filtering process of the standR or generic workflow. Blue contour lines indicate data density. (B) limma-voom mean-variance relationship plots of genes across different biological groups in four GeoMx datasets (see methods). red line: lowess regression, legend as per in A. (C) plot of cell count against total detection for each ROI in the diabetic kidney dataset, colors depicting the distributions of ROIs for individual slide annotation of the ROIs. ROIs with less than 50,000 total detection count (red dotted line) filtered out. (D) After sample filtering, the mean expression-variance plots of gene distribution between standR-filtered or retained samples (right) for either diabetic kidney (top) and colon (bottom) datasets were shown. The linear regression of genes of filtered (blue), retained (green) or unfiltered samples were plotted showing the retained samples maintaining the mean-variance relationship in the data while those samples filtered are different.

For sample filtering: In the *standR* workflow, the distribution of both library size and cell count were taken into account, we used a common library size threshold of 50,000 so that lower-quality ROIs with very small library sizes in the distribution histograms can be removed. To quantify the differences between the original ROIs, filtered ROIs and retained ROIs, we fit a linear model between log-scaled mean expression and the variance of the gene expressions, and the fitted data were then used to calculate the residual sum of squares (RSS).

### Normalization

Different normalisation methods are available in the *standR* package via the *geomxNorm* and *geomxBatchCorrection* functions. When performing RUV4 batch correction, we first identified 200 negative control genes using the *findNCGs* function from the *standR* package. The *geomxBatchCorrection* function was then applied where the parameter k, which indicates the unwanted factor to be used, was set to 3 for the diabetic kidney and brain data, and 2 for the lymph node data. The weight matrices from RUV4 were then included in the design matrix of the linear model as covariates when performing DE analysis.

### Assessing normalisation performance

To calculate the similarity statistics for assessing normalizations, adjusted rand index, jaccard index, mirkin distance and silhouette coefficient were calculated using the *plotClusterEvalStats* function from the standR package between the first two principal components of the data and their slide and tissue type annotations, which indicate batch effect and biological effect, respectively.

### Differential expression analysis

In the generic workflow, paired T-test from the R package *stats* and multiple testing adjustment using Benjamini-Hochberg correction were used to identify statistically significant (FDR < 0.05) differential expressed (DE) genes. In the *standR* workflow, *duplicateCorrelation* from the *limma* R package were first used to calculate the consensus correlation across patients to account for patient variation as a random effect. The linear model was then fitted to the appropriate experimental design containing the biological factors of interest. DE was then performed for specific contrasts of interest, including comparing abnormal glomeruli in diabetic kidney (n=65) to glomeruli in normal kidney (n=12); comparing B cell zone (n=24) to T cell zone (n=24) in lymph node; comparing longitudinal muscle layer (n=8) to circular muscle layer (n=20) in colon; and comparing cortical layer II/III (n=18) to hippocampus CA1 areas (n=13) in brain tissues. The resulting statistic was an empirical Bayes moderated t-statistic, followed by multiple testing adjustment was carried out with the Benjamini–Hochberg procedure to identify statistically significant (FDR < 0.05) DE genes.

### Gene-set over representation analysis

The Molecular Signatures Database (MSigDB) gene-sets (15,16) data was obtained via the R package *msigdbr*. C5 and the Hallmark gene-sets was then used in the over representation analysis. The *enricher* function from the R package *clusterProfiler* (17) was then used to perform the over representation analysis. Gene-sets with adjusted P-value smaller than 0.05 were considered as significantly enriched gene-sets.

## RESULTS

### A comprehensive analysis workflow for Nanstring GeoMx DSP data: standR

From a review of the published studies from Jan 2020 to July 2022, we observed that there is a trend to use the combination of ProbeQC and limitation of quantification (LOQ) filtering strategy to conduct data quality control (QC) (Figure 1B). ProbeQC is the default data processing method provided with GeoMx DSP data, where negative probes are used to detect and remove outliers in the dataset. LOQ is a metric calculated based on the distribution of negative probes and is used as a proxy of the quantifiable limit of gene expression for each tissue fragment (7). After QC, the data is typically scaled using third quantile (Q3) normalization to account for technical variation in the dataset (Figure 1C). Most commonly, differential expression (DE) analysis is performed using standard t-test (Figure 1D). Based on these and for ease of comparison in this study, we define a generic workflow composing these commonly used analysis steps: probeQC and LOQ filtering for data QC, then a Q3 normalisation of the data, followed by identification of differentially expressed genes using a t-test (Figure 1E).

In this study we proposed a refined analysis workflow for Nanostring GeoMx DSP data, which we believe is more suitable for spatial contexture analysis and the complex experimental workflow typically found in Nanostring GeoMx DSP experiments. Here we present the *standR* analysis workflow which consists of recommended strategies for each step (Figure 1F) in a sequential manner.

For QC, our approach is to identify genes that are lowly expressed in over 90% of the regions of interest (ROIs), such genes are then removed from the analysis because genes with constantly low expression are unlikely to be determined as significantly differential expressed genes given their inadequate significance power (18). Subsequently, ROIs with low cell count and/or low total detection count are considered as low-quality tissue fragments and filtered from the analysis to avoid bias due to sample quality in the downstream comparisons.

After QC, suitable normalization method is required due to variation within the Nanostring GeoMx data can be driven by various complex factors, including the desired biological factor such as diseased and control groups or different tissue/cell type groups, or unwanted technical factors such as slide variations (datasets may have each slide containing individual or multiple patient samples), tissue microarray cores differences, different experiment runs or sequencing depth variation (Supplementary figure 1). In such cases where batch effects are observed, it is recommended to apply an appropriate batch effect correction method in the workflow to remove unwanted variation so that fair comparisons between biological groups can be established. Finally, in the standR workflow, DE workflows such as *limma-voom* (11,12) or *edgeR* (13) are preferred instead of standard t-test, as these methods have been shown to be more appropriate for obtaining accurate DE results from complex experimental designs (14).

### Comparison between standR and a generic workflow of commonly used analytical processes

To demonstrate the advantages of using the standR analysis workflow, here we applied both *standR* and generic workflows to analyse four publicly available Nanostring GeoMx DSP datasets from the Spatial Organ Atlas (19), systematically comparing the results generated by the two workflows at different stages of the analyses. The public datasets used are from human diabetic kidney, lymph node, colon and brain tissues, respectively, using the whole transcriptome atlas (WTA) panel for GeoMX DSP (> 18,000 genes).

### standR gene filtering approach retains tissue relevant genes

The basic principles of gene filtering in both workflows are the same: a gene is removed when its expression is smaller than a certain threshold. However, there are three main differences. The first is that the generic workflow uses the distinct LOQ, which is calculated based on the geometric mean of the negative probes measured in the tissue fragments of each ROI separately, while *standR* calculates an overall expression threshold based on both the library size and the minimum count requirement for all genes. The second difference is that the generic workflow uses raw count data while *standR* uses log-transformed count per million (CPM) data to perform the filtering. Comparing the filtering results for all four datasets tested, the generic workflow tends to remove more genes from the analysis than *standR* (Figure 1A and B, supplementary figure 2 and supplementary file 1). In the diabetic kidney and lymph node datasets, *standR* removed markedly fewer genes than the standard filtering, though the genes it did remove were largely also removed by the generic workflow. However, in the other two datasets (colon and brain), the generic removed a substantial proportion of genes, (5.94% and 37.33% respectively), while *standR* did not remove any genes (Supplementary figure 2B). Our comparison also shows that genes filtered by *standR* are outliers across all ROIs for the mean expression-variance distribution while the generic workflow may also remove genes with medium level of mean expression and variance (Figure 2A). For example, in the brain dataset, the generic workflow removed some tissue-relevant genes, such as MDGA1 and CLMP (Supplementary figure 3), which may lead to loss of meaningful biological insight. In particular, CLMP is membrane protein coding gene where the expression of the CLMP gene was reported in the developing cerebral neocortex and other brain areas and might regulate aspects of synapse development and function in the brain (20,21). Similarly, MDGA1 gene encodes a membrane protein, which has a role in cell adhesion, migration, and axon guidance and, in the developing brain, neuronal migration (22,23). We used linear models to investigate if the *standR*-filtered genes are biologically significant (Figure 2B). By taking into account biological factors as covariates in the model, it can be seen that genes filtered by both methods (including *standR*-filtered) are not highly variable between the groups. However, in the brain dataset specifically, there are genes filtered by generic only which are highly variable and potentially DE. Not unexpectedly, MDGA1 and CLMP are amongst them (Supplementary figure 3B labelled), indicating that these two generic-filtered brain-related genes might be differentially expressed between the biological groups in the data.

**Figure 3.**
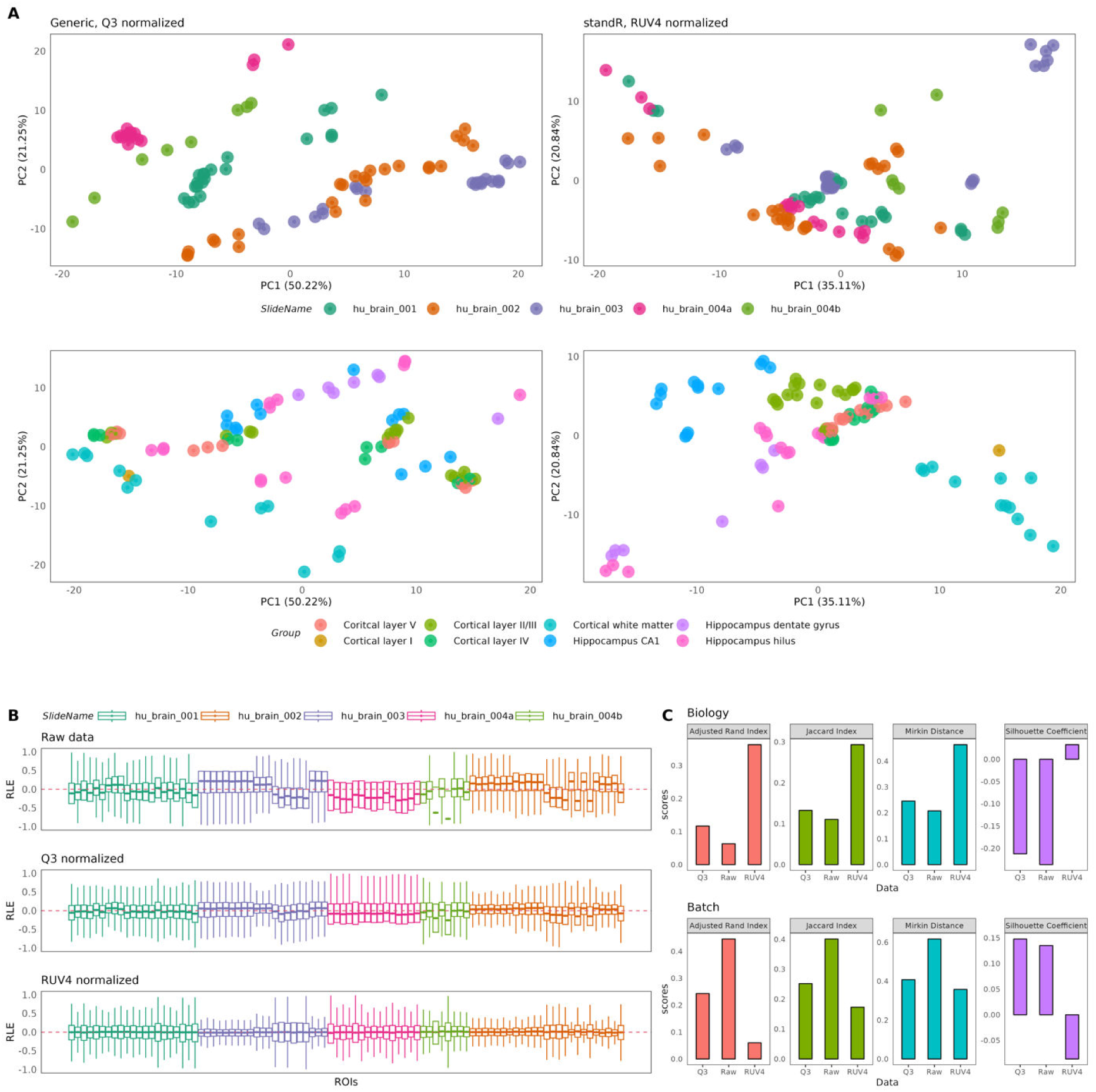
Normalization and batch correction using standR and generic workflows in the brain dataset. (A) PCA plots of data normalised using either the generic (left) or standR (right) workflows. Panels denotes annotations by either ROIs (top) and tissue structures (bottom). standR normalisation and batch correction was able to reduce slide effects while improving separation of biology. (B) RLE plots of the raw (top), Q3-normalised (middle) and RUV4-normalised (bottom) data show RUV4 gives the best removal of technical variations. (C) Summarised statistics of raw, Q3-normalised and RUV4-normalised data comparing performance in terms of the biology (top) or batch factor (bottom). RUV4 performed the best across the statistics in terms of biology and batch (in general).

### standR sample filtering is able to remove low quality samples

During Nanostring GeoMx experiments, low-quality ROIs, such as those with low cell count, might be acquired during the tissue sampling. In order to detect and flag such ROIs, the Nanostring GeoMx NGS pipeline has pre-set cut-offs, such as sequencing read count, sequencing saturation, minimal nuclei count and minimal size of segment area (7,19). As such, the generic workflow does not apply any further sample filtering. However, some low-quality ROIs might not be captured by these pre-defined cut-offs, we therefore included a ROI QC step in the *standR* workflow, which uses the relationship between the cell count and the total detection distribution of each ROI to identify low-quality ROIs (Figure 2C and supplementary figure 4). In this study we applied a common threshold of 50,000 total detection counts to identify low-quality ROIs for the four datasets, removing 8 ROIs in both the diabetic kidney and colon datasets (Supplementary file 2). The mean-variance distribution of the genes for the ROIs that were filtered suggests that genes within them are lowly expressed and less variable compared to the retained ROIs (Figure 2D). Additionally, the residual sum of squares (RSS) (see Methods) between the filtered ROIs and the unfiltered data (103608.2 and 191174.4) is much higher than the RSS between the retained ROIs and the unfiltered data (0.065 and 1.801) (Figure 2D), indicating that *standR* filtered ROIs that are very different from the other ROIs in these two datasets. Taken together, this suggests that the *standR* ROI QC strategy provides an additional filter for low-quality ROIs, supporting a more-accurate downstream analysis.

### Comparison of normalization results

Data normalization can adjust data to a comparable scale by removing undesired biases, such as library size differences, batch variations and other technical factors, allowing a better estimation of the data. In the case of GeoMx data, the experiments are usually composed of multiple slides and patient samples, which can lead to batch effects caused by differences between slides. Furthermore, the heterogeneity and density of cells in the selected ROIs can also lead to variation in library size. Other factors including the age of samples, or sample preparation steps can also introduce variation. It is therefore crucial to perform suitable normalization to allow comparative analysis, such as differential expression analysis, between groups. Technical variations can be visualised in QC plots, such as relative log expression (RLE) plots, which are sensitive to technical variations (24), and principal component analysis (PCA) plots, which visualises the variation in the data by dimension reduction and investigate how these variations are related to the factors in the experiment. In the *standR* package, we provide implementations of different normalization methods, including the “*trimmed mean of the M values”* (TMM) from the *edgeR* (13) and the “*median of ratios”* from *DESeq2* (25), both of which are established data normalization method for bulk RNA-seq data. Similarly, established batch correction methods including *“Removal of Unwanted Variation 4”* (RUV4) (26), “*Remove Unwanted Variation Using Control Genes*” (RUVg) (27) and “*Remove Batch Effect*” function in *limma* (11) are also implemented in *standR*.

PCA was performed on the raw data of the four GeoMx dataset tested. We identified confounding batch effects due to slide differences in the brain, lymph node and diabetic kidney datasets, while no batch effects were identified in the colon (Supplementary figure 5). The generic workflow uses Q3 normalization, which is a method using the 75^th^ quantile as normalised factor for each ROI. In the batch-confounded datasets (i.e., brain, lymph node and diabetic kidney), the PCA of Q3-normalised data suggest that the batch effect due to the different slides has not been removed, nor is the variation explained by tissue types (Figure 3A, left and supplementary figure 6).

Using the *standR* implemented normalisation functions, RUV4 normalization was applied to batch affected datasets (i.e. diabetic kidney, lymph node and brain), while TMM normalization was applied to the colon dataset. Results as shown in Figure 3A (right) suggested improved grouping based on tissue type (bottom, biological) in the brain dataset while reducing the grouping based on slide (top). Results for RUV4 normalisation on diabetic kidney and lymph node datasets are shown in Supplementary figure 6. These suggest that by using appropriate methods provided by *standR*, batch effects can be appropriately addressed. Further evidence of appropriate normalisation outcomes can be found in the RLE plots, which shows less technical variations for standR-normalised data as compared to Q3-normalised data from the generic workflow (Figure 3B). This applies to all four-dataset tested, including the colon dataset, where batch effects are not observed (Supplementary figure 7).

To quantify the performance of the normalization methods, we calculated similarity statistics between the first two principal components of the data and data annotation (Figure 3C and Supplementary figure 8, see Methods). It is clear that normalised data from the standR workflow consistently score high in the statistics comparing biology (i.e. tissue types) and consistently score low when comparing batch (i.e. slide differences). This suggests that application of the appropriate normalization strategy based on the *standR* workflow is able to adjust the data to retain the biological variations in the data, while minimising unwanted technical batch variations.

### Comparison of DE results

DE analysis aims to detect statistically significant genes that are differentially expressed between groups of interest, which are used for the biological interpretation of the GeoMx DSP data and downstream analysis, such as pathway enrichment analysis and network analysis. Instead of applying a traditional paired T-test in the generic workflow to identify DE genes (which assumes that all genes are independent and can be strongly influenced by outliers), the *standR* workflow recommends the *limma-voom* DE pipeline, which borrows information between genes to allows a more precise estimation of biological variation (11,12). Moreover, the *limma-voom* pipeline uses linear modelling which allows greater degrees of freedom and more statistical power and is more useful in the analysis of data from the complex experimental designs typical of GeoMx experiments.

Comparing the DE results between the generic and *standR* workflows, we define one comparison for each of the four datasets: kidney - comparing abnormal glomerulus in diabetic kidney to abnormal glomerulus in normal kidney; lymph node - comparing B cell zone to T cell zone in lymph node; colon - comparing longitudinal muscle layer to circular muscle layer; and brain - comparing cortical layer II/III to hippocampus CA1 areas (Supplementary file 3). We found that the *standR* workflow identified more DE genes than the generic workflow in three of the comparisons, although not in the diabetic kidney dataset (Supplementary figure 9).

By plotting the fold-change and mean expression of genes in a comparison (i.e. MA plots) (28), even dispersion relative to the fold-change are observed in the genes with low average expression in all four comparisons, with the dispersions becoming tighter with higher expression genes (Figure 4A and supplementary figure 10). A skewness of the overall distribution toward negative (in the diabetic kidney data) or positive (in the colon data) log-fold-change can also be seen. This trend is more obvious in the results generated from the generic workflow, suggesting that the log-fold-change is not independent to the expression of the genes, i.e. higher expression comes with higher/lower log-fold-change, which may indicate false positive outcomes.

**Figure 4.**
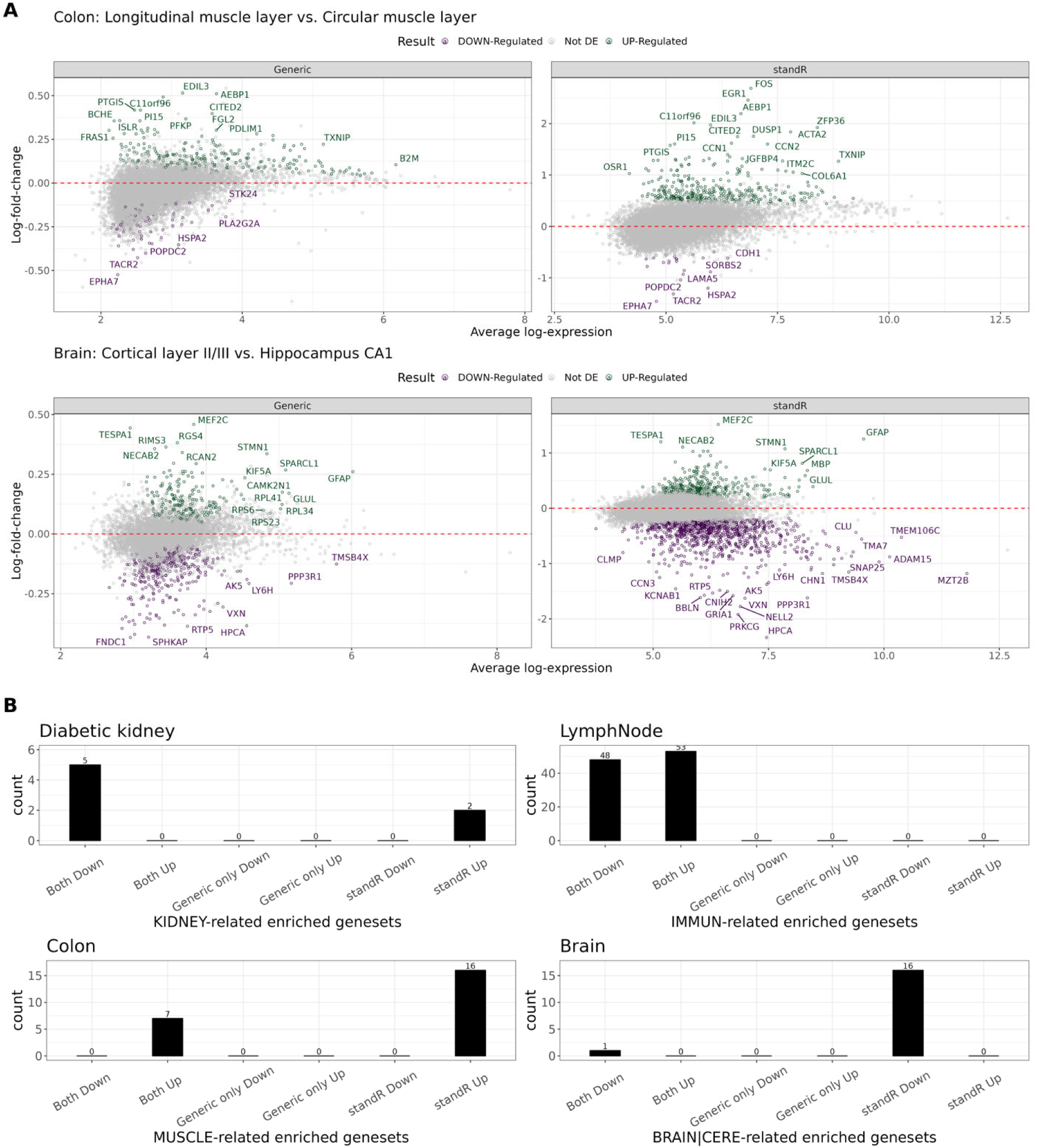
Differential expression analysis results using standR or generic workflows. (A) MA plots visualising differential expressing genes between longitudinal muscle and circular muscle layer in the colon dataset (top) and the comparison between cortical layer II/III and hippocampus CA1 in the brain dataset (bottom) using either generic (left) or standR (right). Differential expression genes generated using the voom-limma pipeline with duplicationCorrelation and applying t-tests relative to a threshold (TREAT) criterion with absolute fold change >1.2 with adjusted p-value <0.05. (B) Gene-set enrichment results of biologically relevant gene-sets were identified for each dataset and the number of biologically-related gene-sets from either workflows were plotted. Overall, standR was able to identify more unique biological gene-sets for the respective dataset.

To assess how well the DE genes identified in either workflow are representative of biological systems, we perform gene-sets over-representation analysis for the up and down-regulated DE genes identified by both or unique to either workflow (Supplementary file 4). In all four comparisons, more biologically relevant gene-sets were found from the DE gene lists identified by either the standR workflow or the intersect between both workflows. For example, in the kidney dataset, kidney-related gene-sets, such as *HP_HORSESHOE_KIDNEY* and *HP_ABNORMAL_LOCALIZATION_OF_KIDNEY,* are found to be enriched in the up-regulated DE genes from the *standR* workflow (Figure 4C). Neither this, nor any kidney relevant results, are found for the generic workflow. Similarly observed in other analysed datasets, the GOBP_MUSCLE_TISSUE_DEVELOPMENT gene-set and the HP_CEREBRAL_CORTICAL_ATROPHY and GOBP_CEREBRAL_CORTEX_DEVELOPMENT gene-sets were only found from DE genes determined by the standR workflow in the colon and brain data analyses respectively. Taken together, this suggests that the *standR* workflow can identify more specific and biologically relevant DE genes. The above observation did not hold for the lymph node dataset, where there was no enrichment for immune-related gene-sets in the DE genes from either workflow, however, there is for the DE genes commonly identified by both workflows (Figure 4B).

## DISCUSSION

### Spatial transcriptomics analysis allows for a greater understanding of the cellular context of disease biology

(29). As one of the key pioneering platforms of this technology, the Nanostring GeoMx DSP offers the ability to study whole-genome spatial transcriptome, with over 18,000 genes for human (22,000 genes for mouse) in a high-throughput manner from both Formalin-Fixed Paraffin-Embedded (FFPE) and fresh frozen materials (7,19). It is crucial to process and analyse the Nanostring GeoMx transcriptome data carefully to identify differential expressed genes with high confidence, leading to a better understanding of spatial transcriptome profiles in the tissues of interest. Here we described *standR*, a Bioconductor package providing quality control, normalization and assessment, and visualization functions for GeoMx transcriptomic data, and recommended a workflow incorporating the well-established *limma-voom* differential expression pipeline to identify DE genes from GeoMx experiments.

There are key issues that differentiates the *standR* workflow from generic workflow, one of which is the gene filtering approach. In the generic workflow, the LOQ was meant for modelling the expression background in each tissue segments to allow removing genes with false signals. However, because LOQ is calculated based on the expression of negative control probes, it will be affected by the cell count and size of each ROI, as well as stickiness or other physical features related to the tissue. In our investigations of the brain dataset, the cell count per segment is negative correlated with the segment area, while the LOQ per segments are positively correlated with the area (Supplementary figure 11). In this case, filtering genes based on LOQ threshold will remove genes with medium expression level and variance (Figure 2A & B, generic only), which may be of biological relevance. This was found in the analysed brain dataset where brain tissue-related genes such as MDGA1 and CLMP were removed by generic only (LOQ). On the other hand, for the standR workflow, the gene filtering threshold is more targeted, using direct calculation based on the expression, while accounting for the library size variation of each ROI. As such, this threshold is relatively stable, and genes with extremely low expression and variation can be accurately detected (Figure 2A&B).

There are currently several Bioconductor packages available analysis of GeoMx spatial data. These include GeoMxTools and GeoMxWorkflows (30,31) which are relevant to Nanostring GeoMx and use linear mixed model as the DE analysis method. The use of the standard paired t-test approach in GeoMx data is inadequate to handle the complexity of the GeoMx datasets. However, in the case of the two new tools, they perform DE analysis using a linear mixed model (LMM)-based method to allow for modelling the individual as a random effect. This is a very useful approach in cross-individual experiments, however their approach is still limited to singular gene comparisons like the paired t-test, which will need large number of replicates in the experiment to increase the degree of freedom and statistical power (32). To address this issue with cross-individual comparisons, the *standR* workflow recommends using the *limma-voom* pipeline with the du*plicateCorrelation* strategy, which not only can borrow information from all genes using an empirical Bayes method to improve the statistical comparison, but can also account for the correlation between individuals by computing consensus correlation between replicates for each gene using restricted maximum likelihood (REML) (33).

The *standR* package and workflow is designed for the analysis of the GeoMx transcriptome data. For GeoMx protein data, the quality control and normalization strategies will be different as the protein panel often contains fewer than 100 markers, with additional housekeeping and IgG markers as negative control markers. Considering the usage of Nanostring GeoMx protein data in the long term and its potential to aid in therapeutics and screening, there is a necessity to develop a comprehensive workflow for analysing both protein and transcript data. With the rapid development of higher plex and finer resolution spatial technologies, specialised analysis workflows and packages, such as *standR*, are essential for ensuring appropriate data QC and processing. While Nanostring GeoMx DSP is reaching maturity as a technology platform, more complex and data rich technologies such as the Nanostring CosMx single molecular imaging platform (CosMx SMI) (34) are now being released. These platforms will also require specialised analysis pipelines and software in order to fully harness the power of spatial location and neighbourhood at the single cell level.

In conclusion, we describe our GeoMx analysis package, standR. We analyse the literature describing GeoMx experiments in order to identify common analytical steps and construct from the most common of these a generic analysis workflow. Then we compare the results from each step between the standR workflow and the generic workflow for four publicly available GeoMx WTA datasets. We provided evidence that standR’s application improves on the detection of biologically meaningful and nuanced results within spatial datasets in comparison to the generic workflow. Overall, we show that the *standR* workflow provides a comprehensive and reasonable quality control process, a better normalization strategy, and a more sophisticated differential expression analysis pipeline.

## AVAILABILITY

Supplementary Data are available at NAR online. The GeoMx DSP datasets used in this paper are available in the Nanostring’s Spatial Organ Atlas (https://nanostring.com/products/geomx-digital-spatial-profiler/spatial-organ-atlas/). The standR package is available in Bioconductor (https://www.bioconductor.org/packages/release/bioc/html/standR.html).

## ACKNOWLEDGEMENT

The authors thank Nanostring Technologies for releasing the publicly available GeoMx datasets, Professor Terry Speed (WEHI) for assistance in the implementation of the RUV-4 function in the standR package and James Monkman and Tony Blick of the University of Queensland for their assistance with reviewing the manuscript.

## SUPPLEMENTARY DATA

**Supplementary file 1. Gene filtering results of GeoMx DSP datasets using generic and standR workflow.**

**Supplementary file 2. Sample filtering results of GeoMx DSP datasets using generic and standR workflow.**

**Supplementary file 3. DE analysis results of GeoMx DSP datasets using generic and standR workflow.**

**Supplementary file 4. Gene-sets enrichment analysis results of GeoMx DSP datasets using generic and standR workflow.**

**Supplementary figure 1.**
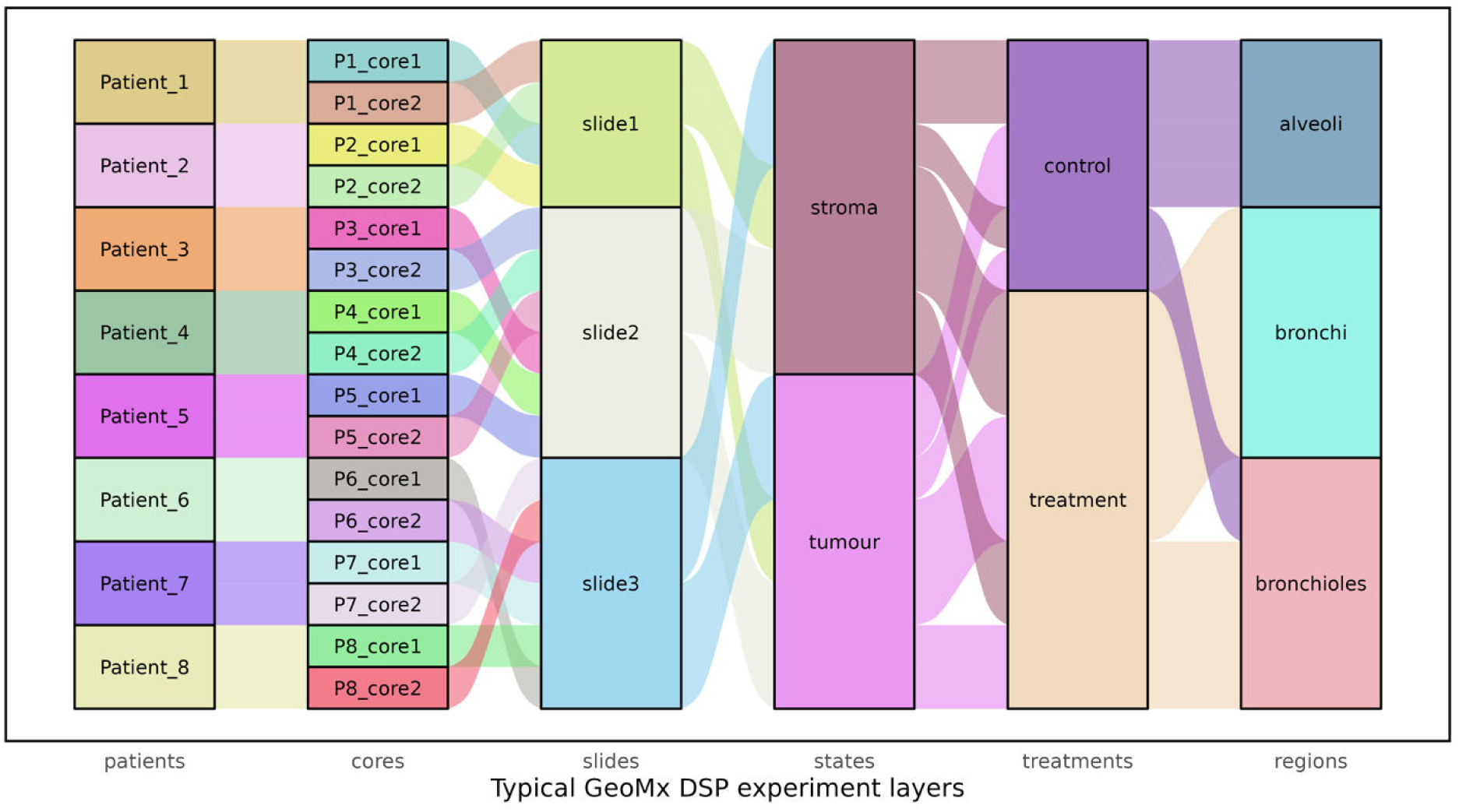
Typical factor layers in a hypothetical GeoMx DSP experiment. Columns are typical biological and technical factors that introduce variations into GeoMx DSP data.

**Supplementary figure 2.**
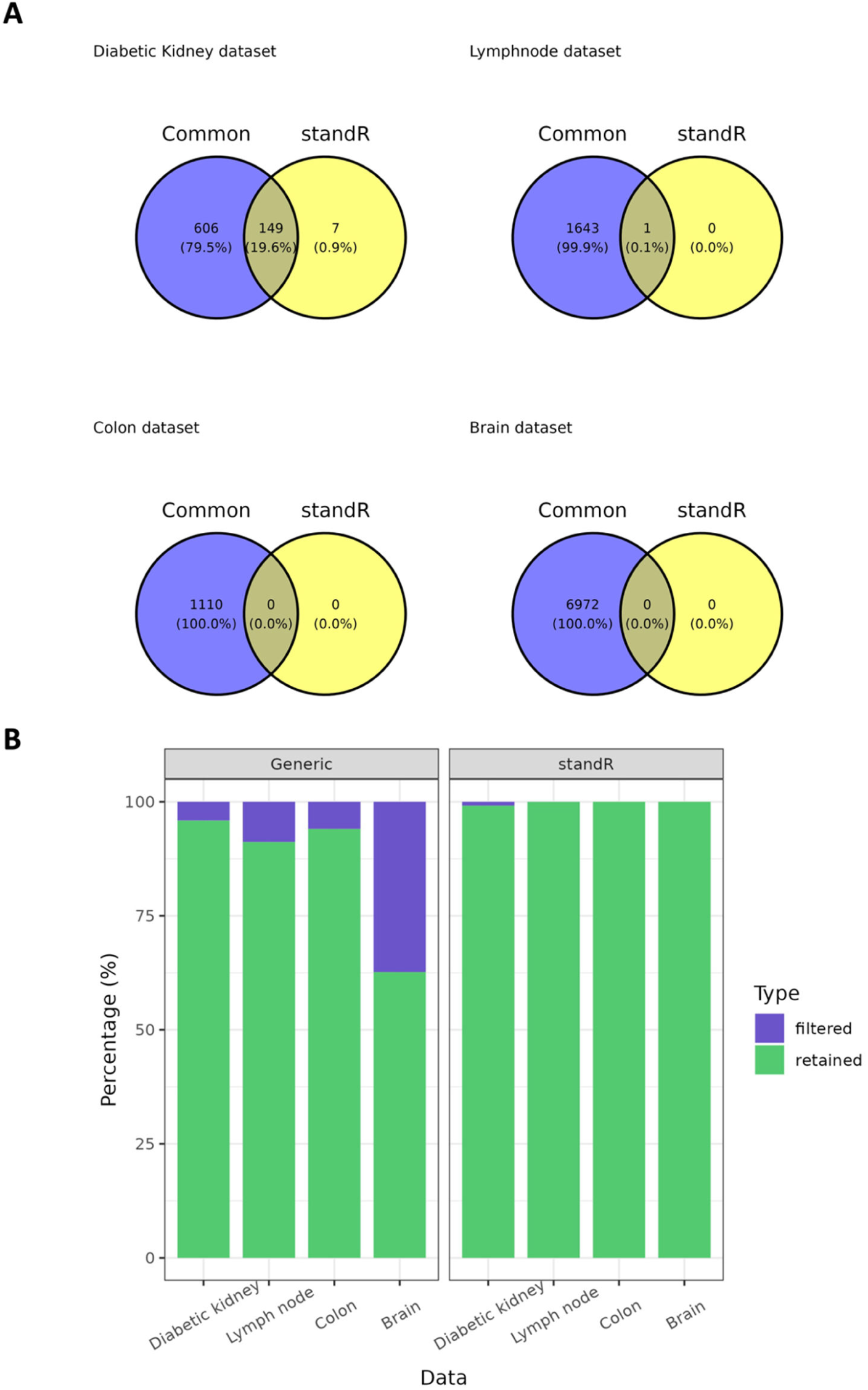
Comparison of filtered genes across four GeoMx datasets. (A) Venn diagrams show the intersection between filtered genes from the generic workflow (purple) and the standR workflow (yellow) for the four GeoMx datasets tested. (B) Percentage of genes either removed or retained during the filtering process of either the generic or standR workflows for the four GeoMx datasets.

**Supplementary figure 3.**
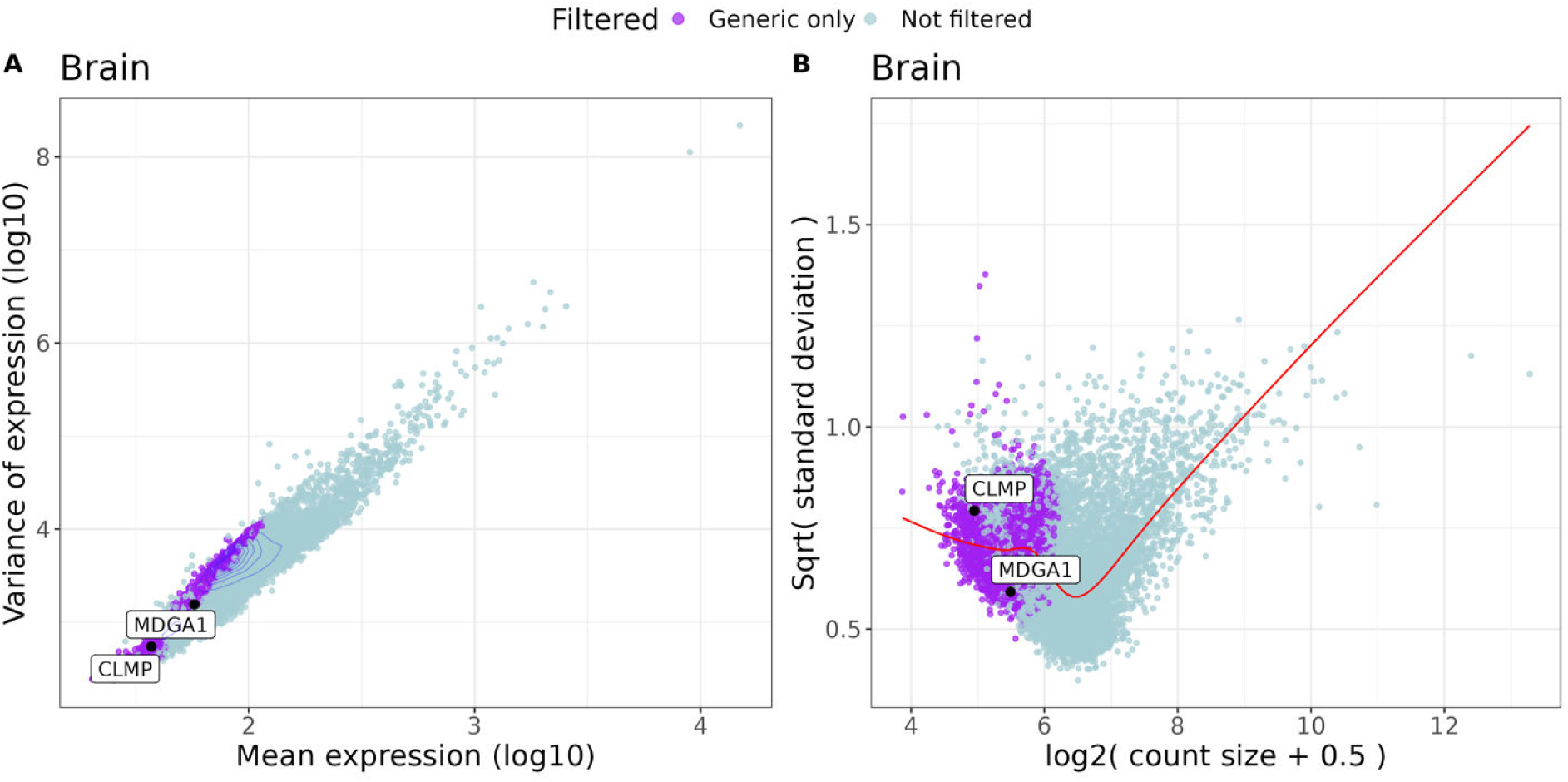
Brain-related genes MDGA1 and CLMP removed by the generic workflow’s filtering. (A) Mean expression against variance of gene expression was plotted for all samples in the brain GeoMx dataset, with purple denoting genes removed by the gene filtering process of the standR or generic workflow. (B) Mean counts (log2) against variance distribution of the genes across different biological groups in the brain GeoMx dataset (see method) was plotted with the red line denoting the lowess regression curve of the data point. Legend colors indicate genes either removed or retained by one or both methods investigated in this study.

**Supplementary figure 4.**
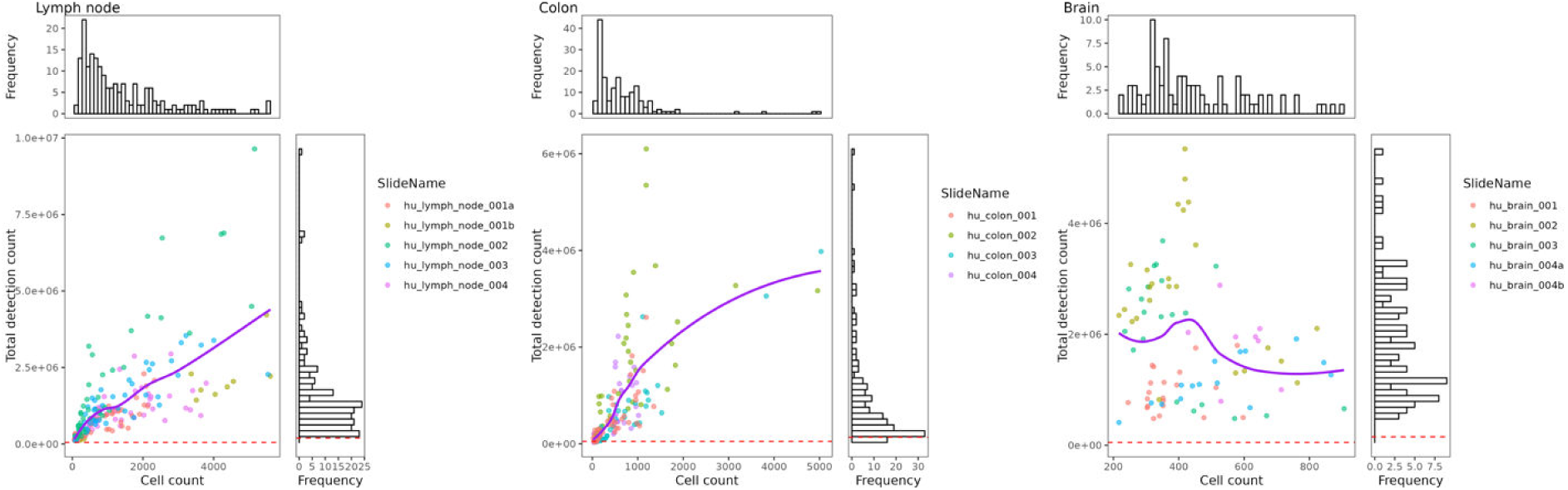
Sample filtering diagnostics and QC in the standR workflow. The cell count vs. total detection of RNA in each ROI of the respective GeoMx dataset was plotted with colours stratifying the ROIs by slide annotations. The purple lines are lowess regression curves with the histograms indicating either the distribution of cell count (top) or total RNA detection (right) of each dataset. ROIs with total detections less than 50,000 (as indicated by the red dotted line) were considered low quality ROIs and removed.

**Supplementary figure 5.**
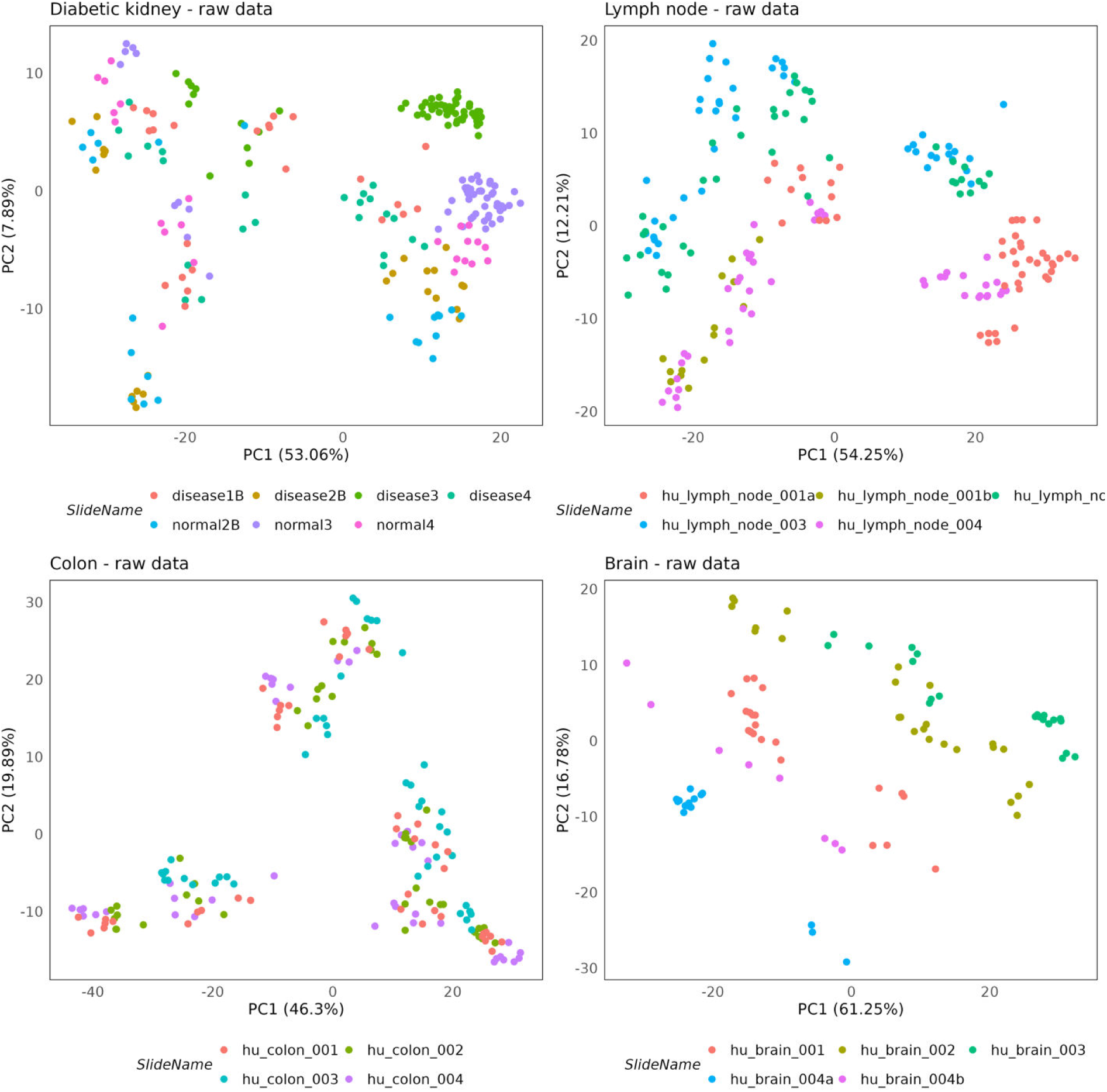
Principal component analysis (PCA) using the raw data for the four GeoMx datasets. The first two Principal Components (i.e. PC1 and PC2) for each dataset are plotted with samples stratified by slide (i.e. SlideName), suggesting that most of the datasets have varying degree of batch effects due to slides, with the only exception being the Colon dataset.

**Supplementary figure 6.**
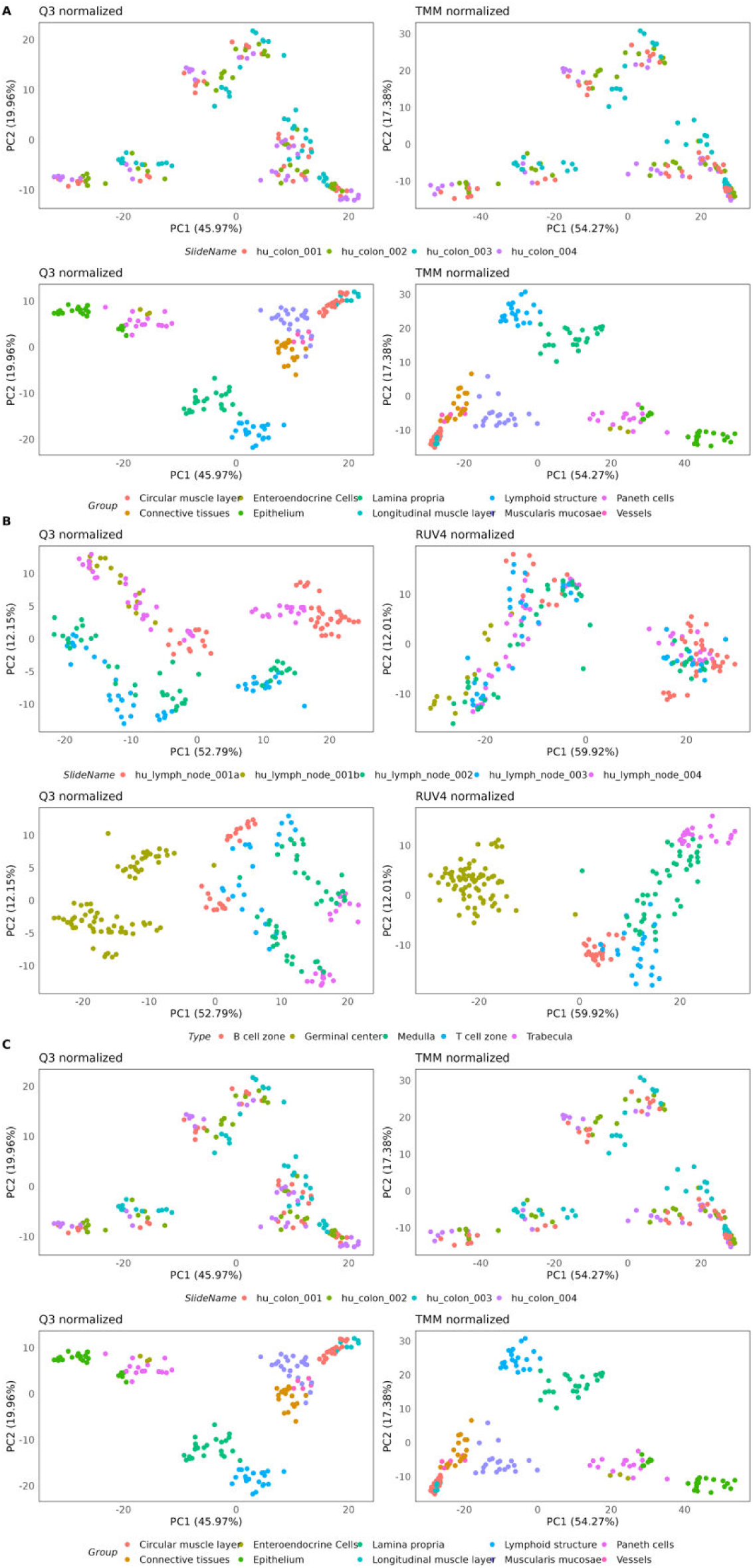
Principal component analysis (PCA) of normalised data for the Kidney (A), Lymph Node (B) and Colon (C) GeoMx datasets. Each dataset consists of 4 panels (2 by 2). The columns on the left are Q3 normalised data generated using the generic workflow, the columns on the right are normalised data generated using the standR workflow. RUV4 was used for kidney and Lymph node datasets, TMM was used for the Colon dataset. For each dataset, top two panels were stratified based on slide while the bottom two panels by tissue types. This shows that standR workflow normalisation removes unwanted batch effects or variations while maintain the biology of interest.

**Supplementary figure 7.**
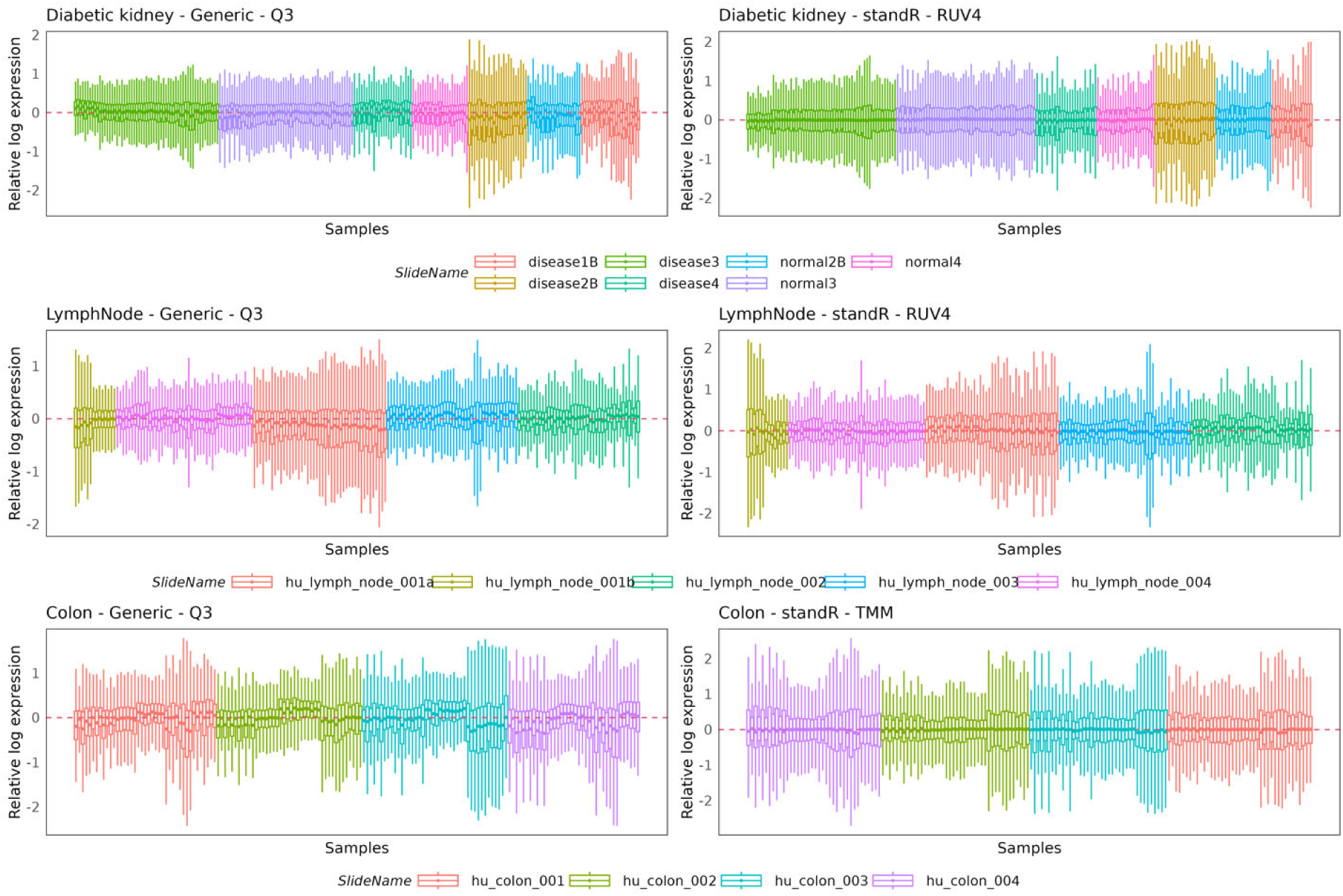
Relative log expression (RLE) analysis of normalised data for the Kidney, Lymph Node and Colon GeoMx datasets. Q3 normalised data generated from the generic workflow are on the left while normalised data generated from the standR workflow are on the right. RUV4 normalisation was used for kidney and Lymph node datasets while TMM was used for the Colon dataset. For each dataset, samples were stratified based on slide. This shows that standR workflow normalisation removes unwanted variations within the respective datasets.

**Supplementary figure 8.**
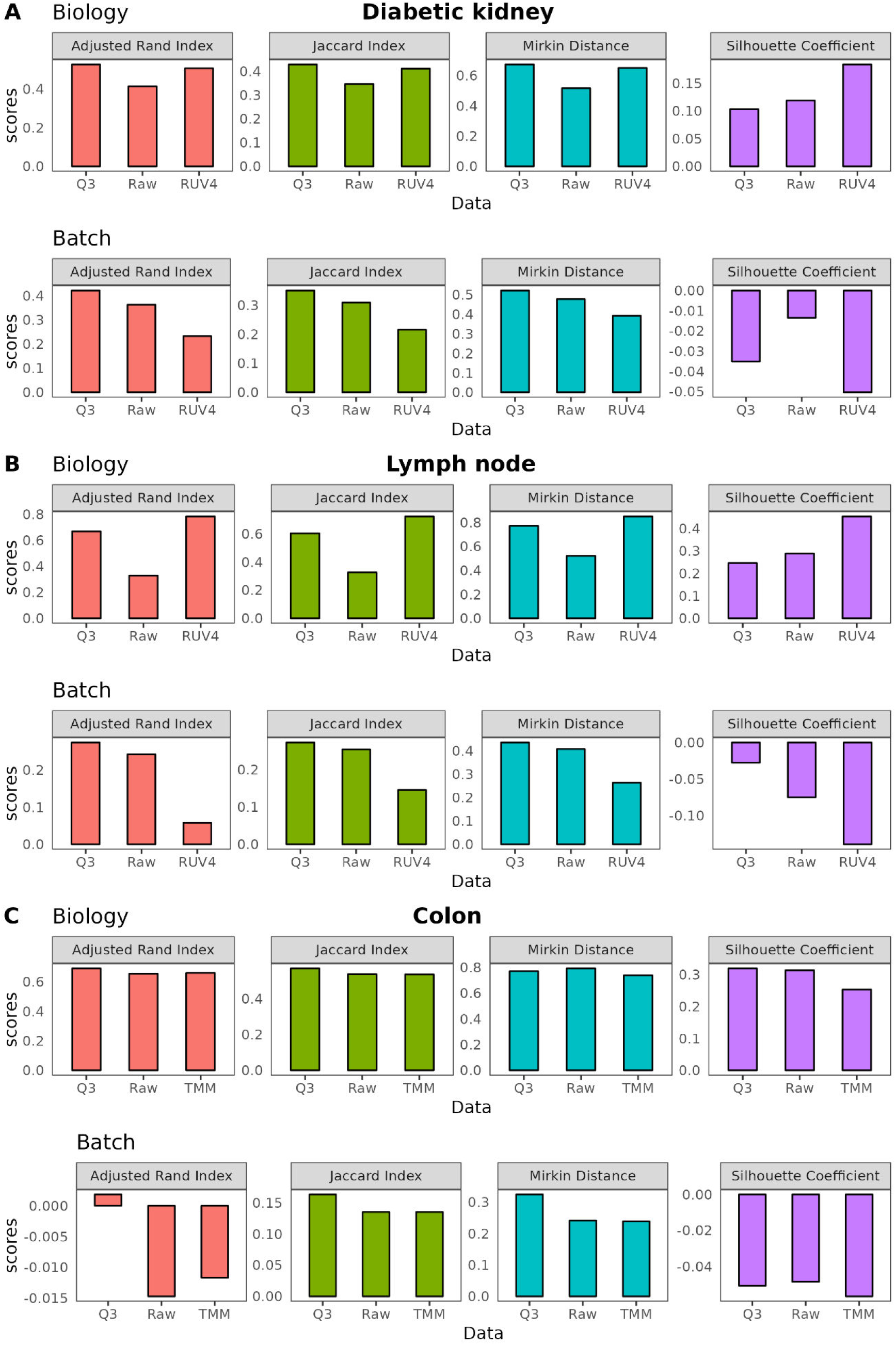
Comparing performances using summarised statistics. Similarity statistics including adjusted rand index, jaccard index, mirkin distance and silhouette coefficient were applied between the first two PCs of the raw, Q3-normalised or RUV4-normalised data of three datasets: (A) diabetic kidney, (B) lymph node and (C) colon, comparing against biology (top) or batch (bottom) as a factor. For (A) and (B), RUV4 consistently scores highly compared to Q3 or unnormalized data when comparing the biology and poorly when comparing batch, suggesting standR workflow (using RUV4) allows appropriate normalisation to be applied to retain the biology while minimising the unwanted technical variations.

**Supplementary figure 9.**
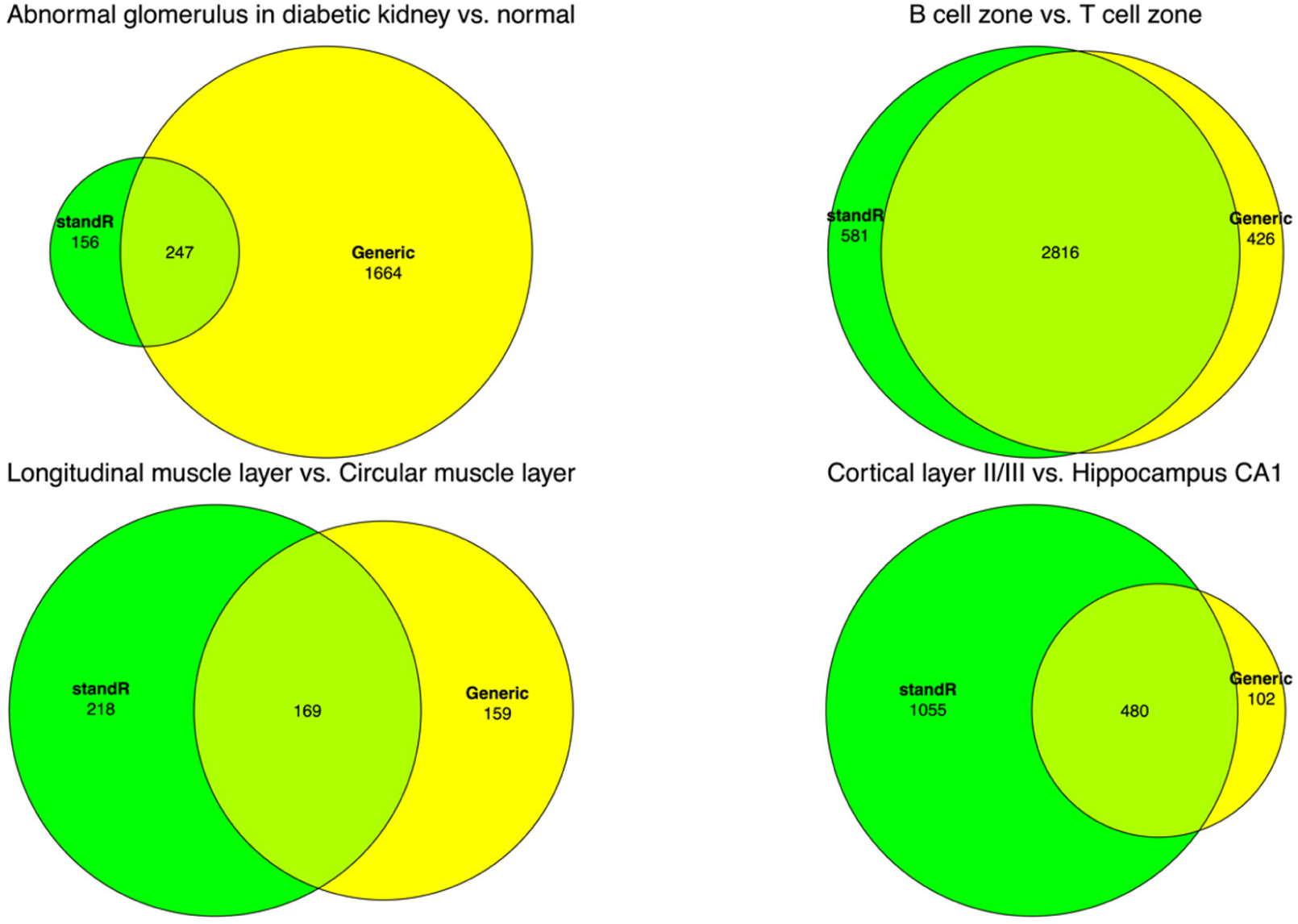
StandR identifies more DE genes in 3 out of the 4 datasets tested compared to the generic workflow. Proportional Venn diagrams of the identified DE genes from either standR or Commonly Used workflows were visualised for each of the 4 tested GeoMX dataset for the respective comparisons applied in this study. Only for the kidney dataset was lesser DE genes identified by standR compared to generic workflow whereas the other three datasets have more DE genes identified by standR.

**Supplementary figure 10.**
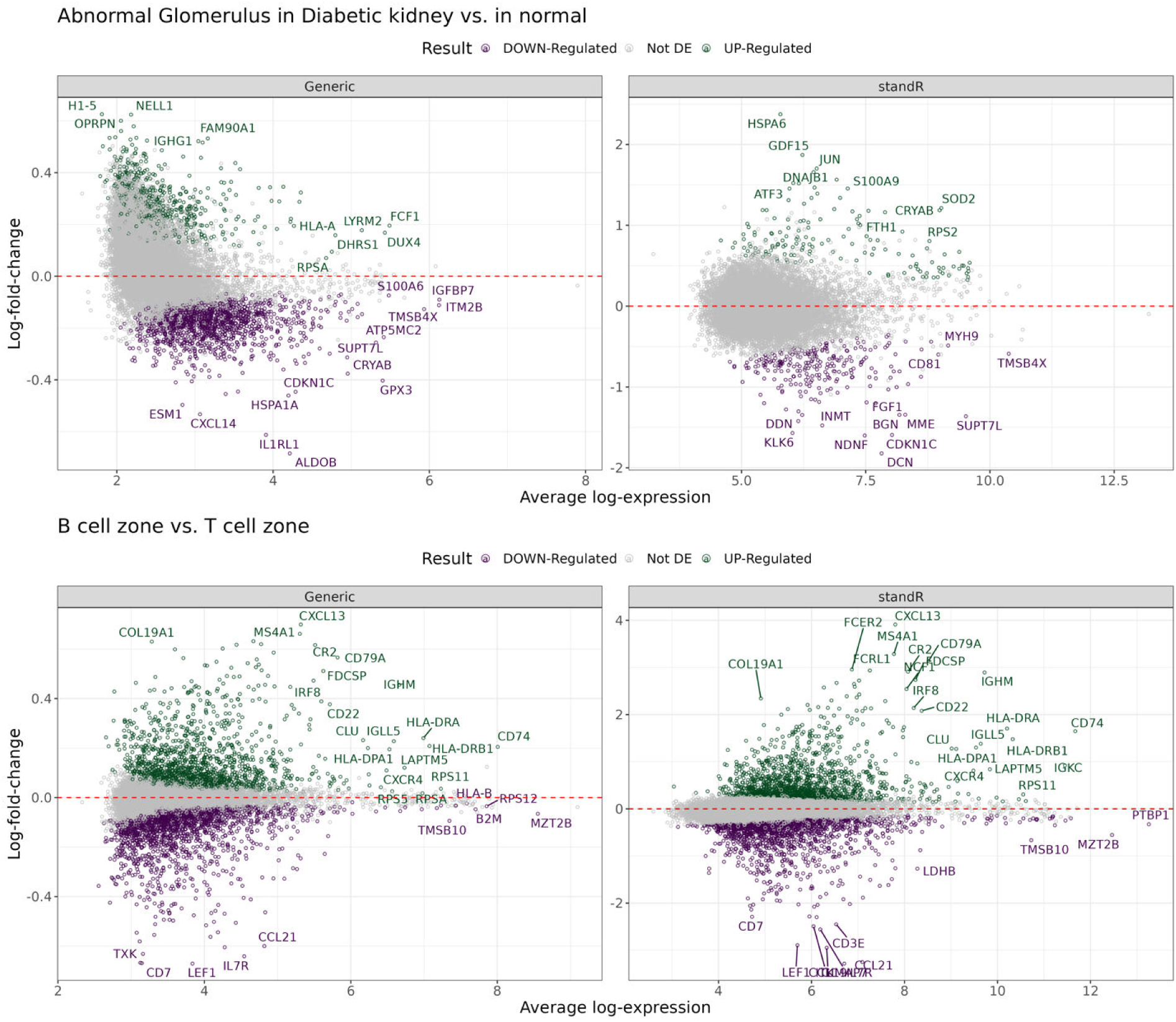
Differential expression analysis results using standR or generic workflows for Kidney and Lymph Node datasets. MA plots visualising differential expressing genes in the comparison between abnormal vs normal glomerulus in diabetic kidney datasets (top) and the comparison between B cell vs T cell zones in the lymph node dataset (bottom). Colours denote significant up- (green) and down-regulated (purple) genes for the respective dataset. Differential expression genes generated using the *voom-limma* pipeline with *duplicationCorrelation* and applying t-tests relative to a threshold (TREAT) criterion with absolute fold change >1.2 with p-value <0.05.

**Supplementary figure 11.**
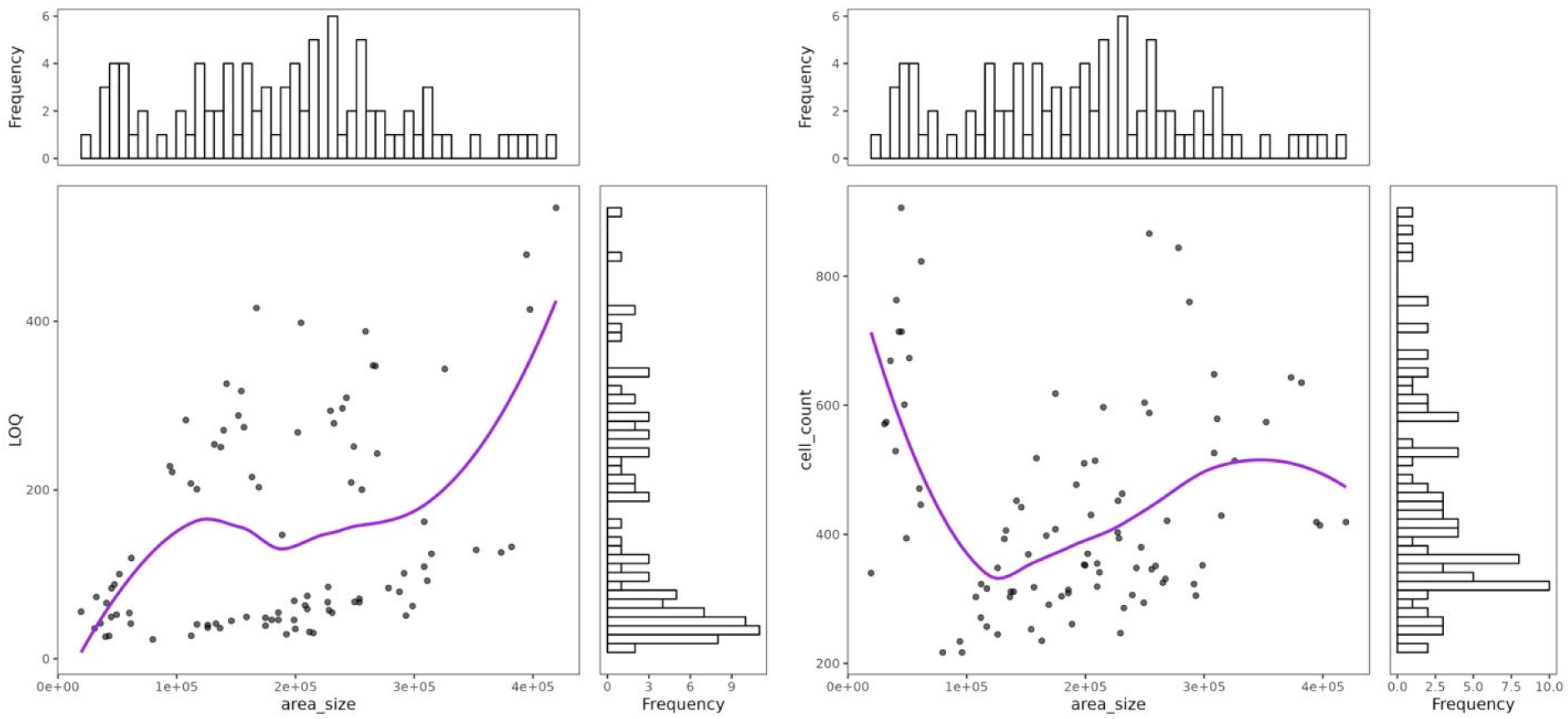
Sample diagnostics and QC using the standR workflow for the brain dataset. (left) The area/size vs LOQ distributions of the ROIs or (right) the area/size vs cell count within the ROIs were plotted for the brain GeoMx dataset. The purple lines are lowess regression curves with the top histograms indicating the distribution by area/size and the right histograms indicating the distribution of either (left) LOQ or (right) cell count of each dataset. ROIs with total detections less than 50,000 (as indicated by the red dotted line) were considered low quality ROIs and removed.

## REFERENCES

1. Cancer Genome Atlas Research, N. (2008) Comprehensive genomic characterization defines human glioblastoma genes and core pathways. Nature, 455, 1061–1068.

2. Ballouz, S., Verleyen, W. and Gillis, J. (2015) Guidance for RNA-seq co-expression network construction and analysis: safety in numbers. Bioinformatics, 31, 2123–2130.

3. Saliba, A.-E., Westermann, A.J., Gorski, S.A. and Vogel, J. (2014) Single-cell RNA-seq: advances and future challenges. Nucleic acids research, 42, 8845–8860.

4. Wu, H., Kirita, Y., Donnelly, E.L. and Humphreys, B.D. (2019) Advantages of single-nucleus over single-cell RNA sequencing of adult kidney: rare cell types and novel cell states revealed in fibrosis. Journal of the American Society of Nephrology, 30, 23–32.

5. Ji, A.L., Rubin, A.J., Thrane, K., Jiang, S., Reynolds, D.L., Meyers, R.M., Guo, M.G., George, B.M., Mollbrink, A., Bergenstrahle, J. et al. (2020) Multimodal Analysis of Composition and Spatial Architecture in Human Squamous Cell Carcinoma. Cell, 182, 1661–1662.

6. Jiang, S., Chan, C.N., Rovira-Clave, X., Chen, H., Bai, Y., Zhu, B., McCaffrey, E., Greenwald, N.F., Liu, C., Barlow, G.L. et al. (2022) Combined protein and nucleic acid imaging reveals virus-dependent B cell and macrophage immunosuppression of tissue microenvironments. Immunity, 55, 1118–1134 e1118.

7. Merritt, C.R., Ong, G.T., Church, S.E., Barker, K., Danaher, P., Geiss, G., Hoang, M., Jung, J., Liang, Y., McKay-Fleisch, J. et al. (2020) Multiplex digital spatial profiling of proteins and RNA in fixed tissue. Nat Biotechnol, 38, 586–599.

8. Moses, L. and Pachter, L. (2022) Museum of spatial transcriptomics. Nat Methods, 19, 534–546.

9. Peixoto, L., Risso, D., Poplawski, S.G., Wimmer, M.E., Speed, T.P., Wood, M.A. and Abel, T. (2015) How data analysis affects power, reproducibility and biological insight of RNA-seq studies in complex datasets. Nucleic Acids Res, 43, 7664–7674.

10. Conesa, A., Madrigal, P., Tarazona, S., Gomez-Cabrero, D., Cervera, A., McPherson, A., Szczesniak, M.W., Gaffney, D.J., Elo, L.L., Zhang, X. et al. (2016) A survey of best practices for RNA-seq data analysis. Genome Biol, 17, 13.

11. Ritchie, M.E., Phipson, B., Wu, D., Hu, Y., Law, C.W., Shi, W. and Smyth, G.K. (2015) limma powers differential expression analyses for RNA-sequencing and microarray studies. Nucleic Acids Res, 43, e47.

12. Law, C.W., Chen, Y., Shi, W. and Smyth, G.K. (2014) voom: Precision weights unlock linear model analysis tools for RNA-seq read counts. Genome Biol, 15, R29.

13. Robinson, M.D., McCarthy, D.J. and Smyth, G.K. (2010) edgeR: a Bioconductor package for differential expression analysis of digital gene expression data. Bioinformatics, 26, 139–140.

14. Seyednasrollah, F., Laiho, A. and Elo, L.L. (2015) Comparison of software packages for detecting differential expression in RNA-seq studies. Brief Bioinform, 16, 59–70.

15. Subramanian, A., Tamayo, P., Mootha, V.K., Mukherjee, S., Ebert, B.L., Gillette, M.A., Paulovich, A., Pomeroy, S.L., Golub, T.R., Lander, E.S. et al. (2005) Gene set enrichment analysis: a knowledge-based approach for interpreting genome-wide expression profiles. Proc Natl Acad Sci U S A, 102, 15545–15550.

16. Liberzon, A., Birger, C., Thorvaldsdottir, H., Ghandi, M., Mesirov, J.P. and Tamayo, P. (2015) The Molecular Signatures Database (MSigDB) hallmark gene set collection. Cell Syst, 1, 417–425.

17. Yu, G., Wang, L.G., Han, Y. and He, Q.Y. (2012) clusterProfiler: an R package for comparing biological themes among gene clusters. OMICS, 16, 284–287.

18. Chen, Y., Lun, A.T. and Smyth, G.K. (2016) From reads to genes to pathways: differential expression analysis of RNA-Seq experiments using Rsubread and the edgeR quasi-likelihood pipeline. F1000Res, 5, 1438.

19. Zollinger, D.R., Lingle, S.E., Sorg, K., Beechem, J.M. and Merritt, C.R. (2020) GeoMx RNA Assay: High Multiplex, Digital, Spatial Analysis of RNA in FFPE Tissue. Methods Mol Biol, 2148, 331–345.

20. Jang, S., Yang, E., Kim, D., Kim, H. and Kim, E. (2020) Clmp Regulates AMPA and Kainate Receptor Responses in the Neonatal Hippocampal CA3 and Kainate Seizure Susceptibility in Mice. Front Synaptic Neurosci, 12, 567075.

21. Chen, A., Liao, S., Cheng, M., Ma, K., Wu, L., Lai, Y., Qiu, X., Yang, J., Xu, J., Hao, S. et al. (2022) Spatiotemporal transcriptomic atlas of mouse organogenesis using DNA nanoball-patterned arrays. Cell, 185, 1777–1792 e1721.

22. Takeuchi, A. and O’Leary, D.D. (2006) Radial migration of superficial layer cortical neurons controlled by novel Ig cell adhesion molecule MDGA1. J Neurosci, 26, 4460–4464.

23. Kim, J., Kim, S., Kim, H., Hwang, I.W., Bae, S., Karki, S., Kim, D., Ogelman, R., Bang, G., Kim, J.Y. et al. (2022) MDGA1 negatively regulates amyloid precursor protein-mediated synapse inhibition in the hippocampus. Proc Natl Acad Sci U S A, 119.

24. Gandolfo, L.C. and Speed, T.P. (2018) RLE plots: Visualizing unwanted variation in high dimensional data. PLoS One, 13, e0191629.

25. Love, M.I., Huber, W. and Anders, S. (2014) Moderated estimation of fold change and dispersion for RNA-seq data with DESeq2. Genome Biol, 15, 550.

26. Gagnon-Bartsch, J.A., Jacob, L. and Speed, T.P. (2013) Removing unwanted variation from high dimensional data with negative controls. Berkeley: Tech Reports from Dep Stat Univ California, 1–112.

27. Risso, D., Ngai, J., Speed, T.P. and Dudoit, S. (2014) Normalization of RNA-seq data using factor analysis of control genes or samples. Nature biotechnology, 32, 896–902.

28. McDermaid, A., Monier, B., Zhao, J., Liu, B. and Ma, Q. (2019) Interpretation of differential gene expression results of RNA-seq data: review and integration. Brief Bioinform, 20, 2044–2054.

29. Marx, V. (2021) Method of the Year: spatially resolved transcriptomics. Nat Methods, 18, 9–14.

30. Ortogero N, Y.Z., Vitancol R, Griswold M, Henderson D. (2022). Bioconductor, Vol. R package version 3.2.0.

31. Reeves J, D.P., Ortogero N, Griswold M, Yang Z, Zimmerman S, Vitancol R, David H. (2022). R package version 1.5.0. ed. Bioconductor 3.16.

32. Smyth, G.K. (2004) Linear models and empirical bayes methods for assessing differential expression in microarray experiments. Stat Appl Genet Mol Biol, 3, Article3.

33. Smyth, G.K., Thorne, N. and Wettenhall, J. (2003) Limma: linear models for microarray data user’s guide. Software manual available from http://www.bioconductor.org.

34. Lewis, Z.R., Phan-Everson, T., Geiss, G., Korukonda, M., Bhatt, R., Brown, C., Dunaway, D., Phan, J., Rosenbloom, A. and Filanoski, B. (2022) Subcellular characterization of over 100 proteins in FFPE tumor biopsies with CosMx Spatial Molecular Imager. Cancer Research, 82, 3878–3878.

